# Bridging Sensory Extraction and Decision Dynamics: Face Pareidolia Emergent from Bottom-Up and Top-Down Interactions

**DOI:** 10.64898/2026.10.23.760570

**Authors:** Hamid Karimi-Rouzbahani, Jason B. Mattingley

## Abstract

Face pareidolia—illusory face perception in everyday objects—offers a unique window into visual ambiguity resolution, yet its reliance on bottom-up sensory versus top-down feedback processing remains debated. Here, we recorded EEG while participants viewed rapid serial visual presentation streams of an object-to-face morph continuum generated using text-to-image diffusion models. Time-resolved representational analyses revealed a spatiotemporal dissociation: early occipitotemporal areas tracked physical features linearly, whereas later frontocentral channels collapsed this gradient into a sigmoidal categorical boundary. Implementing a novel transformation-based representational connectivity analysis, we tracked an initial feedforward transformation of posterior linear traces into anterior sigmoidal representations. Crucially, continuum partitioning demonstrated that unambiguous stimuli were resolved via feedforward flow, whereas ambiguous pareidolic images recruited prolonged top-down feedback. Furthermore, perceptual sensitivity was modulated by animacy priors and directly predicted individual reaction times. Together, these findings establish face pareidolia as a multi-stage recurrent process combining feedforward feature extraction and top-down categorical resolution.

**Significance Statement:** How the human brain transforms ambiguous visual objects into conscious face perception—a phenomenon known as face pareidolia—remains fundamental to understanding perceptual decision-making. By tracking real-time neural activity across a continuous AI-generated morph continuum, this study resolves a longstanding debate between bottom-up sensory and top-down cognitive models of pareidolia. We show that illusory face perception is neither a simple sensory misclassification nor a purely top-down hallucination, but an active, multi-stage recurrent process. Early visual areas rapidly extract physical features, whereas frontal networks convert these signals into categorical decisions, initiating targeted top-down feedback loops specifically when sensory ambiguity is highest. These findings uncover the dynamic neural architecture governing perceptual resolution and provide a quantitative framework for studying visual illusions and clinical perceptual impairments.

## Introduction

The human visual system is remarkably adept at perceiving faces, a capacity critical for social interaction, communication, and survival (Tsao & Livingstone, 2008). This sensitivity is so pronounced that it can sometimes lead to the phenomenon of *face pareidolia*—the illusory perception of face-like structures in inanimate objects (Taubert et al., 2017; see Figure 1A). While it is well-established that pareidolic face images activate core face-processing networks in the brain, the precise cognitive and neural mechanisms responsible for the phenomenon remain debated. Broadly, theoretical perspectives on face pareidolia can be split into two main schools of thought. One assumes face perception is a predominantly sensory, feature-level effect, driven by bottom-up feed-forward mechanisms within occipitotemporal regions including the occipital face area (OFA) and fusiform face area (FFA) (Kanwisher et al., 1997; Rossion et al., 2003). A second view suggests that face pareidolia arises from top-down feedback from frontal brain areas, reflecting cognitive biases, expectations, or rapid template matching that actively constructs a face representation from ambiguous visual inputs (Summerfield et al., 2006; Bar et al., 2006; Liu et al., 2014; Karimi-Rouzbahani et al., 2021).

**Figure 1.**
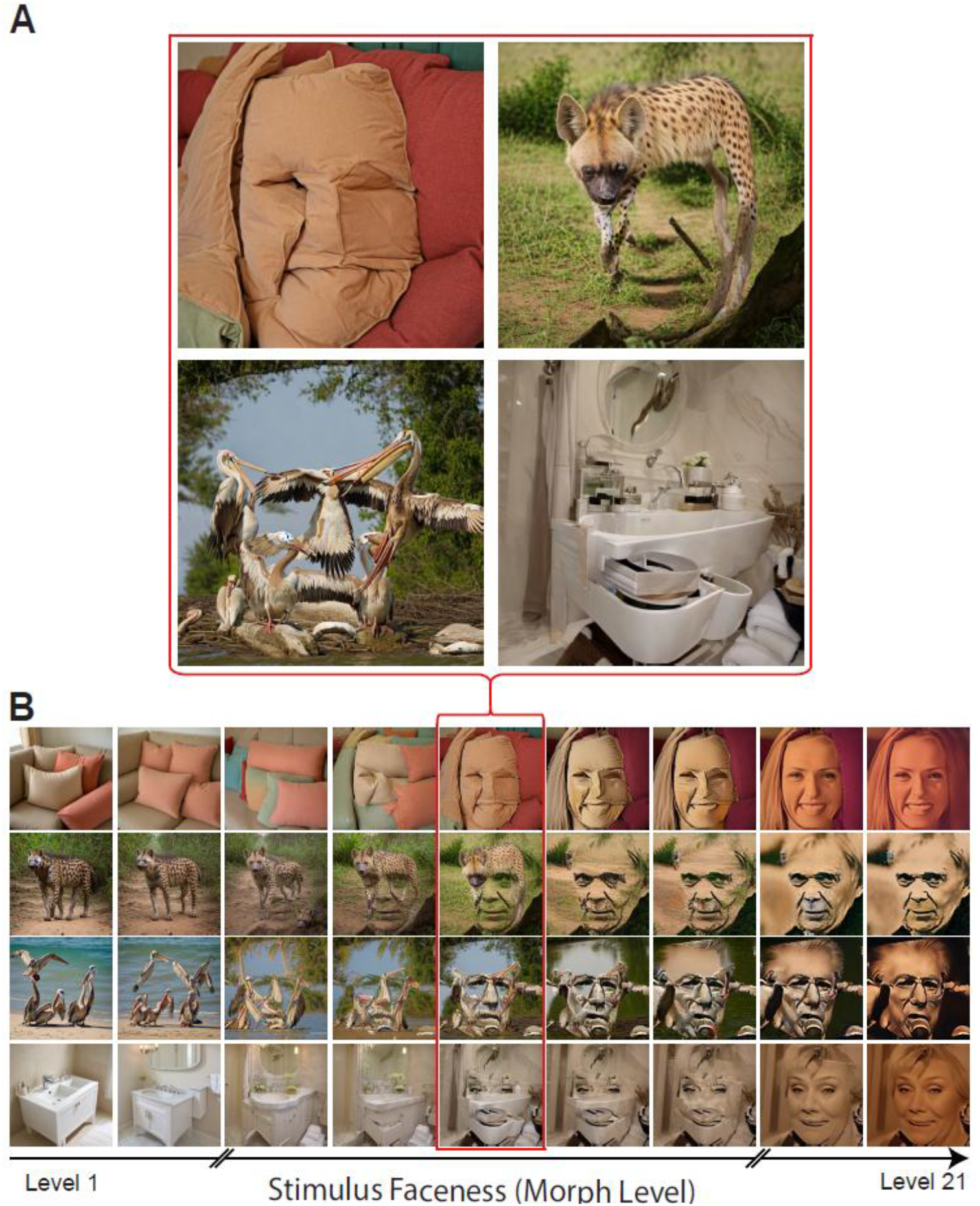
Stimulus design and generative AI morphing pipeline. (A) Mid-level pareidolic stimuli. Representative examples of AI-generated pareidolic stimuli taken from the intermediate stages of the morph continua shown in (B). Face images were obtained from VGGFace2-HQ image set (Chen et al., 2024; https://github.com/NNNNAI/VGGFace2-HQ). These mid-level images illustrate how everyday objects (e.g., cushions, bathroom fixtures) evoke human face perception at ambiguous morph levels. **(B) Generative AI morphing continuum.** Illustrative examples of AI-generated morph continua spanning from unambiguous non-face objects (Level 1) to unambiguous human faces (Level 21). Base images encompassing both inanimate (e.g., cushions, bathroom sinks) and animate (e.g., hyenas, pelicans) categories were morphed through 21 linearly spaced intermediate steps of increasing “faceness”. Intermediate levels represent ambiguous pareidolic stages where visual feature configurations transition from naturalistic object structures to canonical

The perspective that face pareidolia is a low-level sensory effect is grounded in classical models of visual processing, which conceptualise object recognition as a hierarchical, feed-forward cascade (Felleman & Van Essen, 1991; Riesenhuber & Poggio, 1999). In this framework, visual processing is treated as a continuous, linear feature-extraction problem, where visual primitives (e.g., edges, orientations, and spatial frequencies) are extracted in early visual cortex and sequentially integrated into complex configurations within late visual areas (DiCarlo et al., 2012; Kravitz et al., 2013). Because faces possess a highly stereotyped structural arrangement—conventionally described as a T-shaped configuration of two internal horizontal elements above a vertical and horizontal element—visual regions tuned to continuously code these spatial relationships can be inadvertently triggered by matching geometry in inanimate objects (Morton & Johnson, 1991). Neuroimaging work has shown that face-selective areas respond rapidly to pareidolic images (Hadjikhani et al., 2009; Rekow et al., 2022), often indistinguishably from natural faces during initial processing stages (Cerrahoğlu et al., 2025). This rapid and early activation supports the assertion that face pareidolia is a sensory misclassification that arises within the ventral visual stream, operating primarily through proportional, feature-level transformations before higher-order integrative processes intervene (Wardle et al., 2020).

In their search for the earliest sensory evidence for face processing, researchers have investigated selective tuning of the subcortical brain areas—comprising the superior colliculus, pulvinar, and amygdala—which are thought to mediate rapid, non-conscious face detection (Johnson, 2005). Non-invasive electroencephalography (EEG) and magnetoencephalography (MEG) studies have built on these and revealed that face pareidolia manifests as early as 100 to 170 ms post-stimulus, modulating the face-sensitive N170 and M170 components (Churches et al., 2009, 2014). This early temporal window is striking because it matches the timing of natural face processing, and suggests that the structural regularities of pareidolic faces escape initial object-categorisation mechanisms and directly tap into specialised cortical face modules (Wardle et al., 2020). These bottom-up signals appear to be robust enough to carry social cues from inanimate configurations. For instance, behavioural adaptation paradigms have revealed that viewing an illusory (pareidolic) face looking to the left shifts the perceived gaze direction of subsequent real (unambiguous) faces, implying that pareidolic stimuli recruit the same neural populations responsible for coding human social primitives (Palmer & Clifford, 2020). Furthermore, physiological indices such as pupillometry reveal that illusory faces elicit distinct dilation responses that scale with face-likeness, reflecting the persistence of conflicting object and face interpretations rather than a simple intermediate blend (Robinson et al., 2026).

Conversely, a separate body of literature frames face pareidolia not as feedforward extraction of sensory features, but as a discrete categorisation problem driven by top-down modulation and feedback. Rather than passively encoding visual features, frontal areas such as the orbitofrontal cortex (OFC) and ventrolateral prefrontal cortex (vlPFC) are thought to impose categorical decisions onto ambiguous sensory inputs (Bar et al., 2006; Miller & Cohen, 2001; Freedman et al., 2001; Summerfield et al., 2006; Barret & Miller, 2026). According to predictive coding and executive categorisation frameworks, the prefrontal cortex projects an early, coarse prediction about an object’s identity back to sensory areas, effectively collapsing visual ambiguity into a binary or sigmoidal perceptual choice, i.e., assigning the stimulus to either an ‘object’ or ‘face’ category (Friston, 2005; Goldstone & Hendrickson, 2010). In the context of pareidolia, these top-down expectations actively impose a face interpretation onto non-face objects (Lu et al., 2017; Liu et al., 2014; Zimmermann et al., 2019; Palmisano et al., 2023; Romagnano et al., 2024) through a distributed fronto-occipitotemporal network (Karimi-Rouzbahani et al., 2021). Recent MEG and EEG investigations have suggested that while early neural signatures track physical structural dissimilarity, later signals undergo a sharp categorical shift, dynamically transforming from a ‘face-like’ configuration to an ‘object-like’ structure (Robinson et al., 2025; Wardle et al., 2020). These findings imply that face pareidolia involves a dynamic interplay between continuous sensory coding and discrete categorical resolution.

The engagement of such top-down networks is heavily modulated by individual cognitive states, task demands, and underlying neural priors. Neuroimaging evidence indicates that when observers are explicitly primed or instructed to look for faces in noise, frontal and parietal regions exhibit elevated baseline activity, generating predictive top-down signals that artificially increase face selectivity in lower-level sensory areas (Lu et al., 2017) including the fusiform gyrus (Summerfield et al., 2006). This predictive biasing mechanism might explain why face pareidolia is modulated in observers with ‘overactive’ or ‘flexible’ priors, such as people with high levels of paranormal belief or those experiencing clinical symptoms such as hallucinations or Parkinson’s disease, where altered frontotemporal connectivity compromises the validation of bottom-up sensory evidence (Kajiyama et al., 2021; Partos et al., 2016; Powers et al., 2016; Riekki et al., 2013). Furthermore, even among healthy cohorts, substantial interindividual variability exists in how these networks manifest neurally during noise detection tasks (Zimmermann et al., 2019). Functional neuroimaging and connectivity analyses have shown that while unambiguous object images recruit feedforward pathways, illusory face perception is characterised by top-down signals originating in prefrontal structures and modulating ventral temporal areas (Akdeniz et al., 2018; Liu et al., 2014; Romagnano et al., 2024). This top-down imposition suggests that face pareidolia might be a constructive, nonlinear decision process rather than an ‘error’ of the feedforward sensory architecture.

The two theoretical perspectives outlined above make distinct predictions about how neural representations should evolve across gradients of visual ambiguity from unambiguous objects to unambiguous faces, and vice versa. If feed-forward processes are exclusively responsible for face pareidolia, neural representations should track physical stimulus changes in a continuous, linear fashion, reflecting proportional feature-level processing. Conversely, if feedback processes responsible for establishing categorical boundaries in perception are critical, neural representations should exhibit a nonlinear transition—characterised either by a gradual nonlinear sigmoid curve or an abrupt step-function—reflecting the categorical resolution that forces ambiguous stimuli into discrete ‘face’ or ‘object’ categories (Goldstone & Hendrickson, 2010; Philiastides & Sajda, 2006). A limitation of previous investigations is that they relied on static, categorical stimulus designs that compared unambiguous (natural) faces with unambiguous objects and hand-selected pareidolic images (Liu et al., 2014; Wardle et al., 2020). Such binary contrasts cannot arbitrate between linear sensory tracking and nonlinear categorical shifts. Bypassing these limitations requires construction of a continuous, high-dimensional perceptual continuum that bridges ordinary objects and faces.

To achieve this, we leveraged deep generative artificial intelligence (AI) to synthesise a continuous, 21-level morph spectrum that seamlessly transformed ordinary objects into natural faces. Combining EEG with computational representational similarity analysis (Kriegeskorte et al., 2008), we evaluated neural dynamics against three explicit computational models reflecting discrete and competing theoretical mechanisms: (1) a linear model, testing continuous feed-forward feature extraction; (2) a sigmoid model, testing smooth nonlinear categorical boundaries; and (3) a step model testing discrete, binary categorical assignment. We coupled this approach with a transformation-based representational connectivity analysis (Karimi-Rouzbahani et al., 2022) to map directional flow of neural information between posterior sensory and anterior decision-making networks across time.

Foreshadowing the results, we find a clear spatiotemporal dissociation across the brain, such that early posterior areas linearly track the physical progression of visual features across morph levels, while delayed anterior areas collapse this continuous structural information into a discrete, nonlinear sigmoidal representational profile. Crucially, by applying a directed representational flow analysis, we reveal that this architectural transition is mediated by an initial feedforward wave followed by a top-down feedback loop, which is preferentially recruited to resolve categorical uncertainty when sensory signals are highly ambiguous (i.e., pareidolic). The findings suggest that category resolution during illusory face perception involves active, recurrent processing heavily guided by core organisational axes—such as the animate-inanimate continuum—that topologically structure the ventral visual pathway (Mur et al., 2013; Thorat et al., 2019). We also show that the sharpness of these neural decision boundaries is structurally modulated by animacy priors and predicts individual behavioral reaction times.

## Methods

### Theoretical Rationale for Experimental Stimuli and Task Design

To arbitrate between feedforward sensory extraction and top-down categorical decision models of face pareidolia, our EEG paradigm was structured around three key methodological elements: continuous object-face morphing, extended sensory baseline controls, and orthogonal behavioral probe trials. First, pareidolic face perception varies with the level of “face-likeliness” of the stimulus. By spanning a continuous 21-level continuum—along natural faces and synthetic objects—we were able to investigate changes in pareidolic effects especially in intermediate pareidolic morphs (Levels 8–14), which are the most visually ambiguous, and test whether bottom-up sensory and top-down feedback mechanisms are modulated across morph levels. Second, to establish neural baseline anchors and evaluate whether artificial generative morphs engage similar processing pathways to natural images, we incorporated natural object and face images alongside the AI-generated morph continua. These natural stimuli provided independent training sets for multivariate pattern classifiers, allowing us to project the continuous artificial morphs onto a naturalistic decision boundary without circularity. Finally, participants passively viewed the main rapid serial visual presentation streams to isolate unconfounded implicit pareidolia representations. To maintain high attentional engagement without contaminating passive trial EEG epochs with motor artifacts, we dynamically sampled passive probe trials featuring natural faces. Delayed forced-choice gender judgments on these probe trials ensured active vigilance while providing a behavioral reaction-time index to validate neural decision-boundary sharpness.

### Image Set Preparation

To maximise visual control and eliminate low-level visual confounding factors across the continuous morph spectrum, we synthesised our image set using the latent diffusion model Stable Diffusion v1.5 (Rombach et al., 2022), specifically the community fine-tuned checkpoint SG161222/Realistic_Vision_V5.1_noVAE (SG161222, 2023). The latent text-to-image pipeline was governed by three primary parameters: a descriptive textual prompt establishing a baseline object, a target face image, and a morphing scale specifying the progression toward that face (see Figure 1). A linearly spaced 21-level morph sequence was generated using a structured three-step sequential workflow as described below:

1. **Baseline Object Selection:** An object or animal category was selected from the 1000 standard categories defined in the ImageNet Large Scale Visual Recognition Challenge (ILSVRC; Deng et al., 2009), chosen for their widespread use in the field. Examples included categories of *pillow*, *hyena*, *pelican*, and *washbasin*. These categories were then embedded into descriptive text prompts (e.g., *“several pillows on a sofa”*, *“hyena walking in the jungle”*, *“pelicans playing with each other”*, or *“white washbasin in the bathroom”*).
2. **Target Face Selection:** A target face image was selected from the VGGFace2-HQ image set (Chen et al., 2024; https://github.com/NNNNAI/VGGFace2-HQ), which is the processed version of VGGFace2 (Cao et al., 2018), and fed into the model as the endpoint into which the corresponding baseline object category morphed.
3. **Morph Spectrum Generation:** A morphing scale was varied from 0 to 5 in linear increments of 0.25, yielding 21 linearly spaced morph levels for each object-face morph spectrum.

Steps 1 to 3 were repeated to generate 26 distinct object-face morphs/stimulus sets (Figure 1B). Eleven of the object categories morphed from animate objects into faces, and a further 15 morphed from inanimate objects into faces. Sixteen of the target face images were male and 10 were female. In the behavioural experiment, we used this image set which included 546 images (21 morph levels × 26 object-face morph spectra).

For the EEG experiment, in addition to the image sets described above, we also included two natural images of objects (not generated by the diffusion model) and two images of faces per object-face morph. We also included 4 natural face images (2 female) to be used as probe trials in our passive EEG experiment. To minimise the effect of low-level image statistics on neural representations, we converted the images to greyscale via luminance-weighted transformation (rgb2gray in Matlab). Each image matrix was converted to double-precision and normalised by global Z-scoring over the individual pixel profile. Normalised values were then linearly remapped to a standardised [0, 1] intensity range to satisfy image file constraints while preserving internal structural statistics. This resulted in a total of 654 images in the EEG image set (26 object-face morphs × (21 morph levels + 2 natural object images + 2 natural face images) + 4 probe trial face images).

### Ethics Statement

This study was reviewed and formally approved by the Institutional Human Research Ethics Committee at The University of Queensland (2025/HE001279). All procedures were conducted in strict accordance with the Declaration of Helsinki. Prior to participation, all individuals were fully informed of the experimental protocols and provided written informed consent. Participants were explicitly notified of their right to withdraw from the study at any time without penalty, and all data were completely de-identified prior to analysis to ensure confidentiality. Participants were reimbursed for their time at a rate of $20 per hour.

### Behavioural Experiment

To validate the stimulus set and obtain behavioural faceness scores for each image, we conducted a behavioural experiment with 19 adult participants (age range: 21–59 years, mean = 25.1 years; 15 male, 4 female). Stimuli were presented via a computer interface built using the PsychoPy framework.

During the task, the generated pareidolia images were presented in a randomised order to each participant. Each trial consisted of a brief 200 ms stimulus presentation, immediately followed by an 800 ms static Gaussian white noise mask, generated independently at single-pixel resolution on a trial-by-trial basis. Following mask offset, participants reported their subjective perception of the morphed images using a visual analogue slider scale ranging from 0 to 10 (where 0 signified a *complete object* and 10 signified a *pure, human face*). Although responses were untimed to accommodate deliberate judgments, participants were explicitly instructed to register their ratings as rapidly and accurately as possible.

### EEG Experiment

Thirty (30) participants (age range: 19–37 years, mean = 24.5 years; 20 female, 9 male, 1 non-binary/other) completed an electroencephalography (EEG) recording session structured as a rapid serial visual presentation (RSVP) paradigm. The visual display was controlled via MATLAB using Psychphysics Toolbox v3 extensions (Brainard, 1997). Visual stimuli were projected onto a calibrated 1920×1080-pixel LCD monitor positioned at a viewing distance of 60 cm. Stimuli were scaled to subtend a target visual angle of 20deg and centered precisely around the primary coordinate centre x_Centre_, y_Centre_ of the display window.

The experiment was organised into 15 continuous experimental blocks. Each block consisted of an RSVP sequence containing the full set of unique generative morph stimuli, natural face and object stimuli, alongside pseudo-randomly interleaved probe trials (see below). For standard trials, images were presented within the RSVP stream for a strict nominal duration of 50 ms (synchronised to the vertical refresh rate), immediately followed by a nominal 100 ms blank inter-stimulus interval (ISI) containing a central fixation cross (15 × 2 pixels). This rapid timing profile yielded a continuous 150 ms stimulus-to-stimulus onset cycle. Participants were explicitly instructed to passively view the image stream while minimising ocular and muscular movements.

To maintain adequate attention across the extended visual session, 0, 1 or 2 behavioural probe trials were dynamically sampled and pseudo-randomly positioned within every presentation block. Probe trials consisted of the natural probe face images (as described above). When a probe trial occurred, the target face was displayed for a fixed duration of 1.0 s. Following probe stimulus offset, the RSVP stream paused, and an explicit, forced-choice prompt appeared on screen: *“Was the face FEMALE (Left) or MALE (Right)?”*. Participants logged their behavioural responses via a dedicated keyboard interface, using the Left Arrow key for female face judgments and the Right Arrow key for male face judgments. Real-time performance feedback (“Correct!” in green or “Incorrect” in red) was rendered on screen for 1.0 s immediately following key detection. Inter-block intervals allowed participants to rest. Precise timing alignment between the visual presentation display and the continuous EEG recording was achieved via parallel port hardware synchronisation using 10-ms digital pulses.

### EEG Data Acquisition and Preprocessing

EEG data (64-channel) were recorded from participants using a Biosemi EEG system. Preprocessing was performed using MNE-Python according to the protocol described below. Electrode coordinates were mapped to the Biosemi 10-10 channel arrangement across 64 scalp channels. Signals were converted to a common average reference. Digital filtering consisted of a zero-phase band-pass filter from 0.05Hz to 200Hz followed by a 50Hz notch filter to suppress power-line noise. Continuous signals were segmented into single-trial epochs from -100ms to +600ms relative to stimulus onset. Baseline correction was performed by subtracting the mean voltage of the pre-stimulus interval -100ms to 0ms. Artifact-contaminated data segments were mitigated by applying subject-specific adjustments. Specific experiment blocks containing excessive noise were systematically excluded (e.g., blocks 7 and 8 for participant P09, block 1 for P14, block 9 for P15, and blocks 1 through 7 for P23). Clean single-trial epochs were structured into a localised four-dimensional (4D) data array of dimensions: Channels (64) × Time Points × Condition Vectors × Trial Repetitions.

### Topographical Single-Channel Model Fitting

To map the spatial distribution of neural activity, psychometric curve-fitting routines were applied directly to individual EEG channels across 8 target latencies (-100ms to +600ms in 50ms windows around the indicated times). Specifically, for each channel, we built a 21-point response profile (corresponding to the 21 morph levels) by calculating the *mean voltage across repetitions* for each morph level. We then normalised voltage vectors to a relative range [0, 1] via peak-to-peak scaling. Finally, we ran an optimisation algorithm to fit three parameter-constrained functions to the data via nonlinear least squares (Figure 3A): a Linear function (1), a Step function (2) and a Sigmoid function (3)

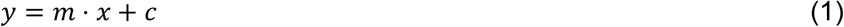

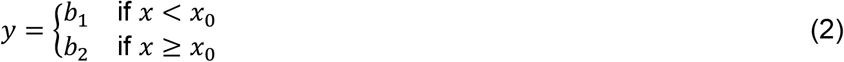

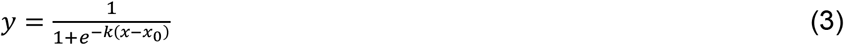

where *c* is the intercept, *x*_0_ is the boundary/inflection point and *k* is the slope parameter. To identify the optimal descriptive representation without parameter-inflation bias, goodness-of-fit was penalised using the Adjusted Coefficient of Determination 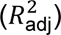. The model yielding the highest positive 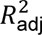 was classified as the “winning model” for that channel.

### Time-Resolved Representational Similarity Analysis (RSA)

To quantify the representational structures of the evolving neural representations, we excuted time-resolved RSA from -100ms to +600ms at 5ms step resolutions.

#### Neural RDM Construction

Neural Representational Dissimilarity Matrices (RDMs) were constructed within each sliding temporal window (5ms). For each time point, clean trial repetitions were split into even and odd indices to facilitate cross-validation. Average spatial patterns across the 21 morph categories were constructed independently for each split. The distance between neural representations was quantified via pairwise Euclidean distances, generating symmetric 21×21 Neural RDMs across two spatial configurations: Anterior Regions (comprising channel Fp, AF, F, FC) and Posterior Regions (comprising channel prefixes CP, P, PO, O).

#### Hypothesis Model RDMs

Vectorised upper-triangular indexing was applied to avoid diagonal symmetry biases. Three mathematical hypothesis profiles were compared against the neural data (see model RDMs in insets of Figure 3A):

1. **Linear Model:** Captures continuous, proportional feature tracking. If the brain tracks physical, low-level visual features of faces across the continuum, its neural representations will exhibit a continuous, monotonic increase across the 21 linearly generated morph levels, modelled using cityblock distance vectors scaled by an inflection point x_0_.
2. **Sigmoid Model:** Captures smooth, nonlinear categorical transitions. If the brain imposes a categorical effect, its neural representations will remain stable at the category edges and show a smooth, nonlinear sigmoidal transition zone at intermediate morph stages, governed by a threshold parameter x_0_and slope gain factor k.
3. **Step Model:** Captures sharp, discrete binary shifts. If the brain behaves as an absolute binary switch (i.e., a face detector), its neural representations will compress into an abrupt mathematical discontinuity, modelled using Hamming distances separating levels below and above a threshold x_0_.

#### Optimisation & Partial Correlation Matrix Mapping

A grid search was performed over the parameter space (*x*_0_ in the range [5, 16]; *k* in [0.1, 0.5, 1.0, 2.0, 5.0]). For each cross-validation fold, hypothesis models were optimised against a training dataset via Spearman rank correlations (rho).

Importantly, to isolate unique variance contributions and remove mutual structural dependencies, optimised models were passed to a partial correlation pipeline using linear regression. For example, to isolate unique Linear coding, the optimised Sigmoid and Step models were designated as nuisance variables. The model RDM and the neural test RDM were residualised against these nuisance models before computing the partial Spearman correlation.

#### Sub-Span Morph Analysis

To examine neural representation profiles in object-like, face-like and ambiguous/pareidolic images, we partitioned the morph spectrum into three categories: ‘objects’ (Morphs 1–7), ambiguous ‘pareidolic’ stimuli (Morphs 8–14), and ‘faces’ (Morphs 15–21). For these restricted sub-spans, we evaluated only the coding (and flow as detailed below) of Linear representations.

### Model-based Representational Connectivity Analysis (RCA)

To trace the dynamics of representational transfer between posterior and anterior brain areas, we performed a novel version of our model-based RCA (Karimi-Rouzbahani et al., 2021, 2022) across temporal inter-areal delays (from 5 to 200 ms in steps of 20ms). This method tracks specific model-defined aspects of information between brain areas. Importantly, it allowed us to remove the effect of unwanted aspects of information on the flow. The pipeline included that, for each target latency t, the historical destination representations (reflected in RDM) at time *t* − Δ*t* were extracted. Instantaneous neural RDM profiles and candidate structural hypothesis models were residualised against this destination history using standard linear regression to isolate novel variance. Partial correlation parameters were recalculated using the history-stripped residuals. We implemented two versions of the method (see details in supplementary Text S1), one looking at transformed representational flow, in which representations turn from a Linear to a Sigmoidal structure, and the other looking at non-transformed representational flow, where representations remain similar across source and destination areas.

- **Transformed** directed flow channels were quantified by taking the product of the historical source representation amplitude and the current destination residual updates:

`o` *Feedforward Transformed Flow (FF):* Calculated as Posterior Linear representations at *t* − Δ*t* multiplied by Anterior Sigmoidal representation at *t*.
`o` *Feedback Transformed Flow (FB):* Calculated as Anterior Sigmoidal representations at *t* − Δ*t* multiplied by Posterior Linear representation at *t*.
- **Non-transformed** flow control pathways were tracked concurrently by isolating equivalent model updates between identical representation structures (e.g., Linear → Linear and Sigmoid → Sigmoid).

To evaluate the capacity of RCA to detect transformations of representations across regions, we simulated time-resolved multivariate pattern data for two interconnected areas (A and B) using a 4×4 factorial design (16 conditions). Area A encoded spatial position during an early window, which was transformed into behavioural category information in Area B after a transmission delay (Δ*t* = 6 time points; see more details about our simulation in Figure S1 and Karimi-Rouzbahani et al., 2022). As predicted, while non-transformed RCA (i.e., Position → Position and Category → Category) failed to detect represenational flow due to the altered representational format, transformed RCA successfully detected a robust, time-resolved peak in representational flow aligned with the simulated (known) transmission delay.

### Time-Resolved Multivariate Pattern Analysis (MVPA)

To establish the generalisability of the AI-generated imageset to naturally captured images (those captured from real-world objects and faces), a multivariate pattern analysis (MVPA) decoding pipeline was implemented. Specifically, a linear classification architecture was constructed using a StandardScaler normalization stage paired with a Linear Support Vector Classifier (LinearSVC, maximum iterations = 5000). The classifier was trained using clear, naturally captured object and face images, and tested on the independent AI-generated morph sequences. Category evidence was quantified by calculating the mean distance of the test patterns to the object-face decision boundary. MVPA was run across the -100ms to +600ms window sampled at discrete 5ms increments.

### Brain-Behaviour Correlations

To assess behavioural relevance, neural properties were correlated with task performance. The four-parameter sigmoidal fitting routine was applied to each channel during a target window centred at 250ms (±150ms window width) for each individual block. The highest slope gain parameter (k) observed across the channel array was extracted as the peak neural slope index for that block. Behavioural task performance for the corresponding block was quantified by calculating the mean reaction time (RT) for correct probe trials. The relationship between block-level neural slope values and behavioural RTs was evaluated using Pearson product-moment correlation coefficients (*r*).

## Statistical Analyses

Group-level significance testing across information coding and representational flow dynamics was conducted using one-sample *t*-tests evaluated against a null hypothesis of zero baseline-corrected effect (*H*_0_: *μ* = 0), as well as paired-sample *t*-tests for cross-condition comparisons. To evaluate statistical strength continuous with effect magnitude, objective evidence tracking was performed using the Jeffrey-Zellner-Siow (JZS) Bayes Factor (*BF*_10_) with a narrow Cauchy prior scale of *r* = 0.5, which places higher prior probability on small-to-moderate effect sizes typical of time-resolved neural data (Morey & Rouder, 2011; Rouder et al., 2009).

Effects and cross-condition differences were classified as statistically reliable when meeting an evidence threshold of *BF*_10_ > 3. For individual condition traces, time segments meeting threshold with positive directional amplitudes (*μ* > 0) were highlighted with increased stroke width. For cross-condition pairwise comparisons, continuous significant intervals (*BF*_10_ > 3) were indicated by horizontal marker bars below the waveforms and the corresponding Bayes Factor traces underneath. To maintain visual readability across figures, time-resolved *BF*_10_ sequences were smoothed using an *n*-point convolutional moving-average window and upper-bounded at 1 × 10^10^ to prevent floating-point overflow.

## Results

### 1. Nonlinear perception across linear morphs

Our generative image set was designed under the prediction that linear increments of object-to-face morphs would yield a gradual increase in face-like perception. As predicted, participants showed a gradual increase in their reports of “faceness” (Figure 2), consistent with previous studies reporting that pareidolic images are perceived at intermediate levels of face-likeness, i.e., more face-like than unambiguous objects and less face-like than unambiguous faces (Wardle, et al., 2020; Robinson et al., 2025). Interestingly, however, there was a clear departure from linearity in participants’ judgements. On average, participants reported a low, unchanging perception of faceness across the first 4 morph levels (i.e., object-like images) followed by a sharper increase in faceness in morph levels 5 - 9, and then a more moderate slope thereafter. These results suggest that although stimuli were generated using linearly spaced morphing, participants’ perception of faces exhibited distinct nonlinearities. There was also a degree of variability in faceness reports across participants (Figure 2, light grey lines).

**Figure 2.**
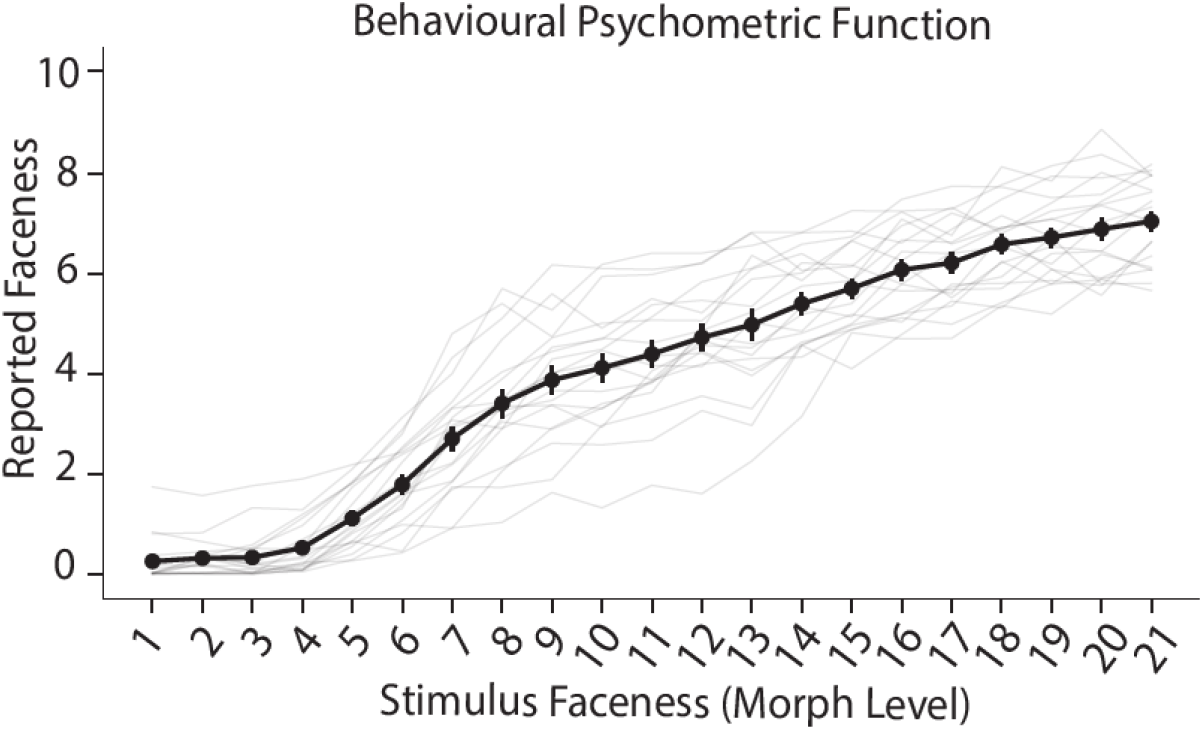
Behavioural psychometric function across the object–face morph continuum. Perceived “faceness” ratings plotted as a function of stimulus morph level (Levels 1–21). The solid black line represents the group-level mean psychometric curve (±SE), revealing a clear nonlinear, sigmoidal response profile despite the linearly spaced morphing of the stimuli. Individual participant responses are depicted by light grey lines, highlighting inter-individual variability in categorical sensitivity and the specific morph threshold at which pareidolic face perception emerges.

### 2. Anterior scalp regions encode sigmoidal category boundaries

Analysis of the behavioural data confirmed a nonlinear sensitivity to face perception along the object–face continuum, despite the linearly spaced morphing of stimulus components. This raises the question of where and how the transition from object to face perception occurs in the brain. To address this, we formalised three hypotheses regarding the neural mechanisms underlying the transformation of linearly spaced physical inputs into nonlinear perceptual decisions (Figure 3A):

1. **Linear Representations:** If an area of the brain tracks physical, low-level visual features of faces, its neural representations will exhibit a continuous, monotonic and linear increase across the 21 linearly generated morph levels.
2. **Sigmoid Representations:** If an area of the brain reflects a categorical effect, its neural representations will remain stable at the category edges and show a nonlinear sigmoidal transition zone at intermediate morph stages.
3. **Step Representations:** If an area of the brain behaves as an absolute, discrete binary switch (i.e., a face detector), its neural representations will compress into an abrupt discontinuity.

To evaluate whether and how these candidate strategies were implemented in the brain, we fitted Linear, Sigmoid, and Step functions to the neural representation profiles across individual electrodes within specific temporal windows. We evaluated and compared the goodness-of-fit across these functions using cross-validation with the adjusted coefficient of determination 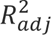.

**Figure 3.**
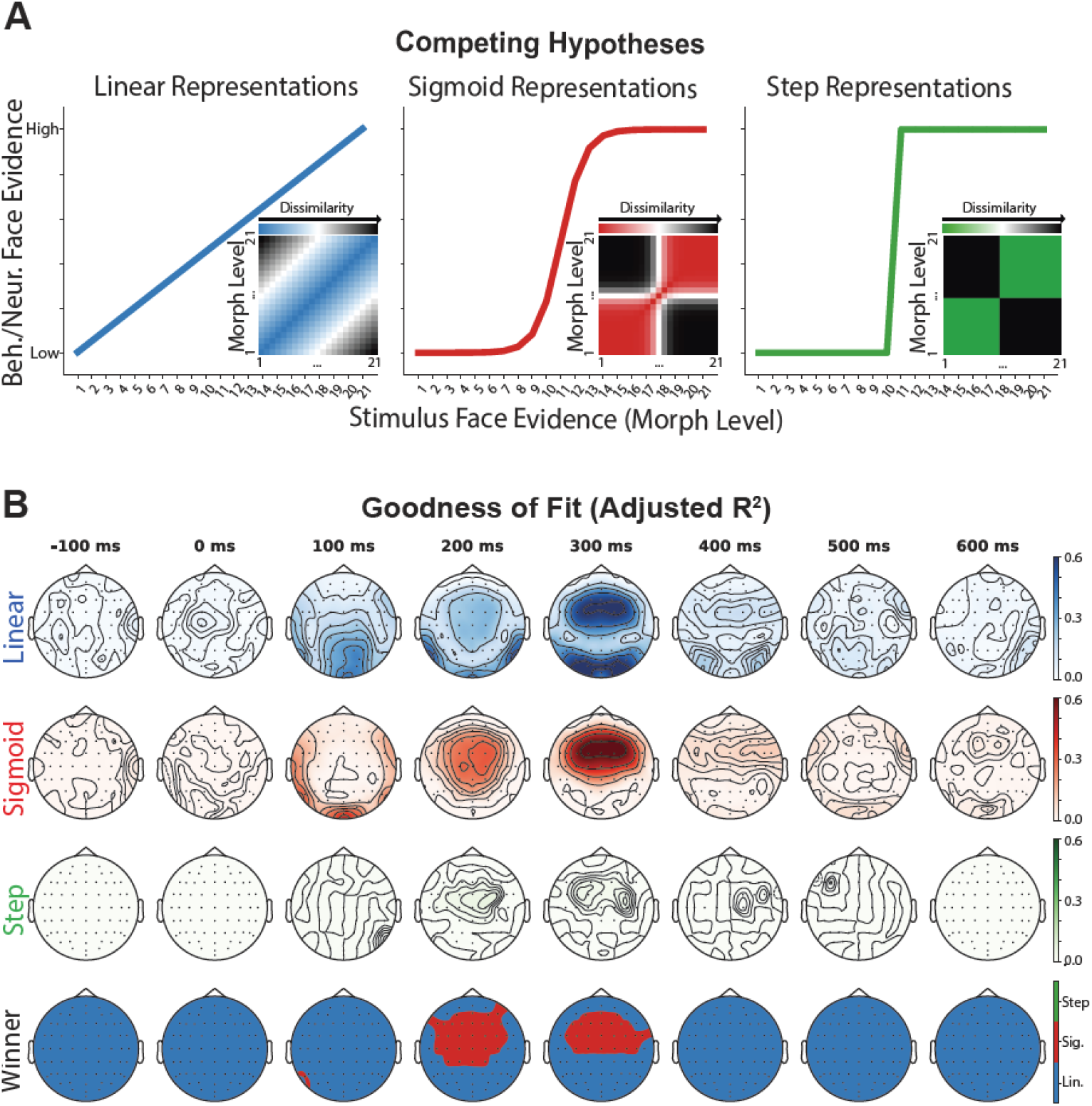
Candidate representational models and spatiotemporal model fits. **(A) Competing representational hypotheses.** Theoretical models evaluating candidate profiles of information representation across the 21 morph levels, alongside their corresponding representational dissimilarity matrices (RDMs; insets). The **Linear** model (blue) hypothesises continuous, proportional tracking of physical stimulus changes. The **Sigmoid** model (red) posits a smooth, nonlinear categorical transition with intermediate ambiguity. The **Step** model (green) predicts a abrupt, binary threshold boundary separating objects from faces. **(B) Spatiotemporal goodness of fit (adjusted** *R*^2^**) and winning models.** Topographical maps depicting model fits (adjusted *R*^2^) across peri-stimulus time points (-100 ms to 600 ms) for each candidate profile across scalp EEG channels. The top three rows display spatial distributions of explained variance for Linear, Sigmoid, and Step models, respectively. The topomaps with empty distributions indicate unsuccessful fits to the Step function. The bottom row (“Winner”) indicates the winning representational model at each channel and time point based on maximal goodness of fit.

Linear representations outperformed the other two functions across most electrodes and time windows (Figure 3B). However, Sigmoid representations outperformed the alternative functions in the 200 ms and 300 ms time windows, specifically across central and anterior regions of the headmap. Step representations never dominated over Linear or Sigmoid representations. These results indicate that, outside of centro-frontal areas, the rest of the brain—including posterior sensory regions—represent the object-to-face morph continuum linearly, reflecting low-level physical visual features of the stimuli. Frontal regions, however, represent the stimuli categorically. This reflects their involvement in making the final categorical decision, potentially pushing the percept of ambiguous (pareidolic) images toward a discrete object or face category.

### 3. Spatiotemporal dissociation of feature tracking and category resolution

Building on the univariate differences observed between posterior and anterior regions across the object–face morph continuum, we next applied multivariate representational similarity analysis (RSA) to evaluate the temporal dynamics of information coding against our three operational models (Figure 3A). Multi-channel activity patterns were analysed in anterior and posterior regions independently. To isolate the unique variance explained by a specific representational profile (e.g., Linear), the contributions of the alternative candidate models (e.g., Sigmoid and Step) were controlled for using partial correlation. Finally, to ensure that models with greater parametric flexibility (e.g., Sigmoid relative to Linear) did not gain an unearned advantage during fitting, we implemented a 2-fold cross-validation scheme by partitioning the dataset into odd and even trials.

Results revealed a clear spatiotemporal dissociation in how the brain processes the object–face continuum, which was heavily dependent on scalp location and post-stimulus latency (Figure 4A). In posterior areas, there was evidence (BF > 3) for a Linear representation of the morph continuum during early time windows, which surpassed Sigmoid and Step representations and peaked at approximately 100 ms. This initial Linear dominance was followed by a peak in Sigmoid representations at approximately 200 ms. At around 300 ms, there was evidence (BF > 3) that Linear representations once again surpassed the alternative models (Linear versus Sigmoid and Linear versus Step). Overall, there was insufficient evidence for Step-like representations in posterior areas.

**Figure 4.**
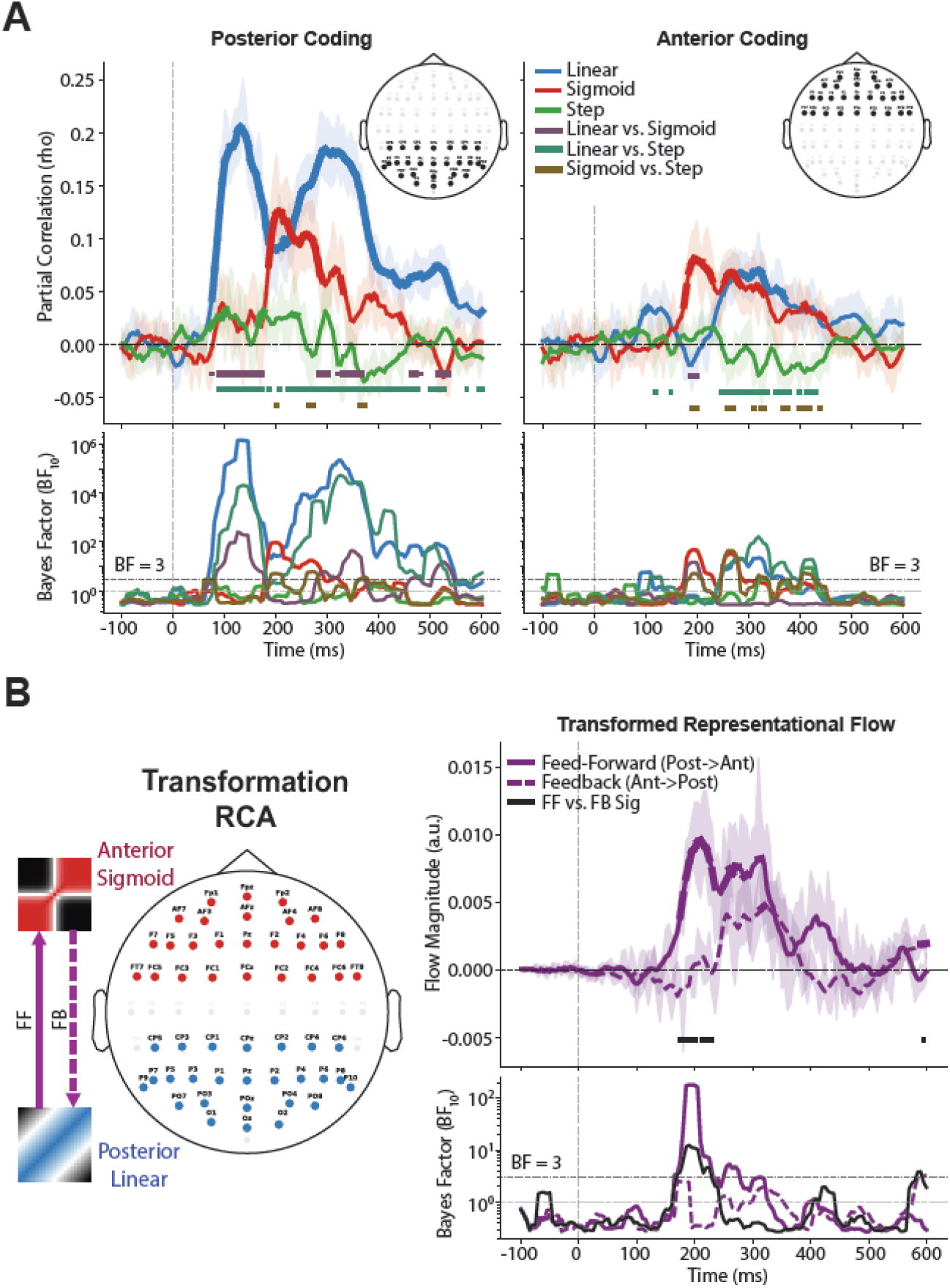
Spatiotemporal representational dynamics and inter-areal information transformation. **(A) Time-resolved representational models in posterior and anterior ROI channels.** Partial correlation trajectories (rho; top panels) and corresponding Bayes Factors (BF_10_; bottom panels) for candidate models (Linear, Sigmoid, Step) across peri-stimulus time in posterior (left) and anterior (right) channel groups (insets). Horizontal bars denote temporal clusters with evidence (BF > 3) for pairwise model comparisons. **(B) Transformed representational flow (Transformation RCA).** Inter-areal representational connectivity tracking the transformation from posterior Linear representations to anterior Sigmoid representations (left schematic). Feed-forward flow (posterior-to-anterior; solid purple line) and feedback flow (anterior-to-posterior; dashed purple line) magnitudes are plotted alongside difference trajectories (black line; top right panel) with corresponding Bayes Factors (BF_10_; bottom right panel). Horizontal black bars indicate time points over which there was evidence (BF > 3) for higher feed-forward than feedback flows.

In anterior areas, by contrast, there was evidence (BF > 3) for brief Linear representations at around 100 ms, followed by evidence (BF > 3) for Sigmoidal representations at around 200 ms that surpassed Linear and Step representations. This Sigmoidal profile persisted until Linear representations reappeared at around and after 300 ms. As in posterior areas, there was insufficient evidence for Step-like representations in anterior regions. These multivariate information-based RSA results confirm the topographic univariate results (c.f., Figure 3), demonstrating that posterior brain areas initially (∼100 ms) encode continuous structural representations of physical stimulus changes without discretising ambiguous sensory inputs—aligning with their role in early visual feature extraction. Conversely, categorical decision-related information is predominantly represented later (from 200 ms onward) in anterior brain areas, establishing the Sigmoid model as the winning strategy for frontal categorical resolution.

### 4. Transformed connectivity reveals feedforward-to-feedback loops

We found that posterior areas represented the object–face continuum in a predominantly linear fashion, whereas anterior areas exhibited a sigmoidal profile. It is unclear, however, whether Sigmoidal representations in anterior regions were generated independently or if Linear representations in posterior regions were transferred and transformed into Sigmoidal representations in anterior areas to support categorisation. To address this, we developed, validated (Figure S1), and implemented a novel transformation-RCA framework to evaluate inter-areal exchange and representational transformation. Importantly, because RCA incorporates the concept of Granger causality, it controls for the influence of past within-area representations on current patterns, isolating transferred information while minimising confounding effects from local, independent processing. Given that Step representations yielded no significant effects, they were excluded from subsequent analyses for brevity.

RCA revealed a sequential bidirectional loop (Figure 4B). There was evidence (BF > 3) for feedforward representational flow from posterior to anterior regions starting at 170 ms and peaking at 200 ms. This representational flow was specifically characterised by posterior Linear representations directly driving and transforming into anterior Sigmoid representations. There was evidence (BF > 3) that these feedforward flows surpassed the subsequent feedback flows (note evidence (BF > 3) for feedback between 320-340ms)—configured here to test whether anterior Sigmoidal representations project backward to modulate posterior Linear representations (for which there was insufficient evidence overall; BF < 3). Under predictive coding and hierarchical processing frameworks, such feedback is expected to project higher-order categorical hypotheses (sigmoid states) downward to modulate or constrain lower-level feature tracking (linear states). These results suggest that representational transformations occur predominantly in the feedforward direction, translating image-level structural profiles into categorical decisions.

To ensure that the transformed flow dynamics reported above could not have been captured by non-transformed Linear-to-Linear or Sigmoid-to-Sigmoid flows, we applied the same RCA while keeping the target representations in source and destination areas identical (either Linear or Sigmoid; Figure S2). In contrast to the transformed flows, which were highly phase-locked to stimulus onset and sequential, non-transformed flows—which tracked communication strictly within an un-transformed model (e.g., Linear-to-Linear or Sigmoid-to-Sigmoid)—showed only a delayed, brief appearance at 265 ms (BF > 3), with weaker evidence for feed-forward flow (compare BF = 8 for Linear-to-Linear here with BF > 100 for Linear-to-Sigmoid transformed flows in Figure 4B). Evidence for purely Sigmoid-to-Sigmoid, non-transformed flows was insufficient (BF < 3). These results support the conclusion that the primary operational link between posterior and anterior brain areas during the perception of our object–face image set involves a structural translation from linear feature tracking in posterior regions to categorisation in anterior regions.

### 5. Pareidolic ambiguity selectively recruits top-down feedback

The findings presented so far distinguish the role of posterior areas in tracking continuous structural representations from that of anterior areas in extracting categorical representations. To test how representational flow supports structural-categorical transformation for pareidolic images specifically, we partitioned the 21 morph levels into three distinct subsets: Object-like (Levels 1–7), Pareidolic (Levels 8–14), and Face-like (Levels 15–21) stimuli. To evaluate how these subsets are represented across regions, we applied sub-span analyses by restricting the linear model (Figure 3A) to these specific morph ranges and computing partial correlations against the empirical neural data. This approach allowed us to test whether directional communication is invariant across object–face morph levels or if top-down feedback is modulated (and potentially enhanced) to resolve categorical uncertainty for pareidolic images. Based on prior work using RCA which showed that perceptual ambiguity enhances feedback flow of information during face recognition (Karimi-Rouzbahani et al., 2021) and visual categorisation (Karimi-Rouzbahani et al., 2026), we predicted that pareidolic images would be represented in a noisy manner in early sensory areas, thereby triggering feedback flows to resolve categorical uncertainty.

Time-resolved analyses across morph levels revealed that the strength and temporal dynamics of information coding and flow were profoundly modulated by categorical uncertainty (Figure 5A). For images at the extreme ends of the spectrum (i.e., Object-like and Face-like), the brain achieved rapid, efficient resolution. In posterior areas, there was evidence (BF > 3) that these relatively unambiguous stimuli evoked strong coding starting at around 100 ms and peaking at approximately 205 ms. In anterior areas, information coding for these stimuli was comparatively weaker and delayed (Figure 5A). By contrast, for Ambiguous Pareidolic Images—where object and face features overlapped maximally—a distinct neural profile emerged. In posterior areas, there was evidence (BF > 3) for lower (nearly halved at around 200 ms) and delayed (peaking at 220 ms) coding for Pareidolic images relative to Object- and Face-like images. In anterior areas, the coding of Pareidolic images was even further delayed relative to Object- and Face-like images, peaking at 330 ms (c.f., 210 ms for Object- and Face-like images). These results suggest that while the brain robustly and rapidly encodes clear semantic categories of objects and faces, the processing of Pareidolic images is significantly protracted and attenuated across both posterior sensory and anterior cognitive networks.

**Figure 5.**
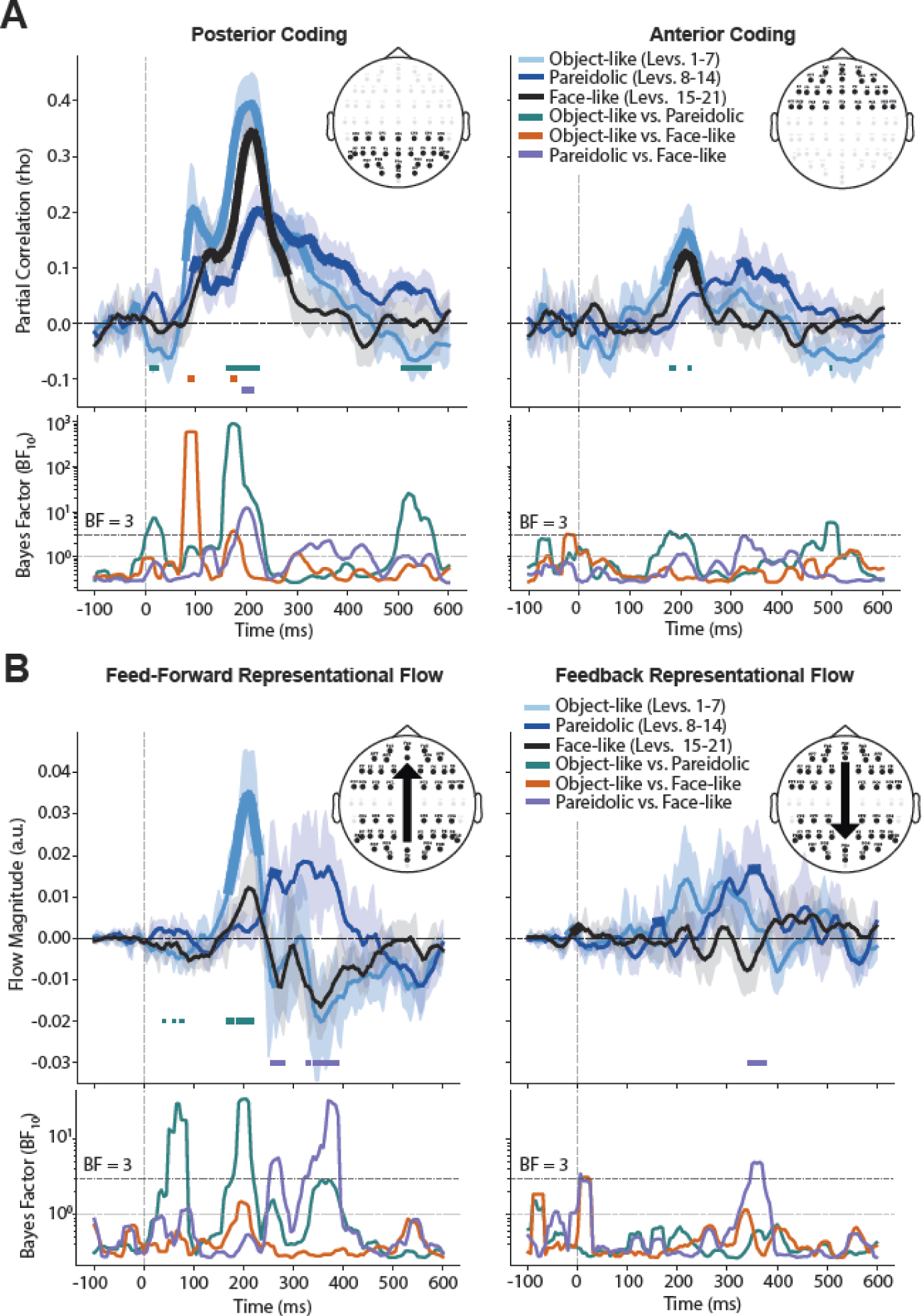
Impact of categorical uncertainty on representational coding and inter-areal information flow. **(A) Time-resolved information coding across morph subsets.** Partial correlation trajectories (rho; upper panels) and corresponding Bayes Factors (BF_10_; lower panels) for Object-like (Levels 1–7; light blue), Ambiguous Pareidolic (Levels 8–14; dark blue), and Face-like (Levels 15–21; black) stimuli across peri-stimulus time in posterior (left) and anterior (right) ROI channels. Coloured horizontal bars indicate time windows with evidence (BF > 3) for pairwise category comparisons. **(B) Category-specific directional representational flow.** Feed-forward (posterior-to-anterior; left panels) and feedback (anterior-to-posterior; right panels) flow magnitudes across time for the three morph subsets, alongside corresponding Bayes Factors (BF_10_; lower panels). Horizontal bars denote time points over which there was evidence (BF > 3) for pairwise flow differences.

To determine whether the top-down anterior to posterior flow of information was modulated by the categorical uncertainty of Pareidolic images, we quantified representational flow between these regions (Figure 5B). Consistent with the information coding results, there was a weak and delayed feed-forward representational flow for Pareidolic images compared with Object-and Face-like images. Specifically, there was evidence (BF > 3) for feed-forward flow of Object-like representations starting at 170 ms, surpassing Pareidolic flows (BF > 3) and peaking at 210 ms. For Pareidolic stimuli, conversely, there was evidence (BF > 3) for feed-forward flow only briefly around 270 ms, showing a delayed peak at approximately 330 ms. Interestingly, while there was no evidence for feed-forward flow of Face-like images, there was evidence (BF > 3) for stronger feed-forward flow for Pareidolic than for Face-like images across several time points.

The delayed feed-forward flow for Pareidolic images relative to Object-like images, alongside the emergence of feed-forward flow for Pareidolic images but not clearer Face-like images, supports the purported engagement of frontal networks to resolve categorical uncertainty. This mechanism was further substantiated by stronger anterior-to-posterior feedback flows for Pareidolic images compared with Object- and Face-like images in the 300-400ms window. Specifically, there was evidence (BF > 3) for feedback of representations exclusively for Pareidolic images at around 170 ms which peaked at 370 ms, with evidence (BF > 3) confirming stronger feedback for Pareidolic than for clearer Face-like images. Feedback flows for Pareidolic images remained active over an extended duration, providing a direct neural correlate of prefrontal recruitment during heightened categorical uncertainty. It is noteworthy that while information coding for pareidolic stimuli was overall attenuated relative to the two unambiguous categories, the preferential recruitment of enhanced feedback flows for these ambiguous images is non-trivial, underscoring a key functional role for top-down processing in resolving category uncertainty.

### 6. Animacy priors and neural boundary sharpness predict behavior

We next evaluated whether our AI-generated morphed stimuli evoked neural representations comparable to naturalistic object and face images typically used in the pareidolia literature (e.g., Romagnano et al., 2024; Robinson et al., 2025). This was critical to ensure that AI-generated stimuli did not induce idiosyncratic representations outside the established object–face processing pathways. To test this, we trained whole-brain multivariate pattern classifiers on EEG responses to the naturalistic (unambiguous) object and face images included in the experiment (see Methods) and projected the AI-generated morphed stimuli onto the learned decision boundary (Figure 6A). We then calculated the decision-boundary distance for each morphed image.

**Figure 6.**
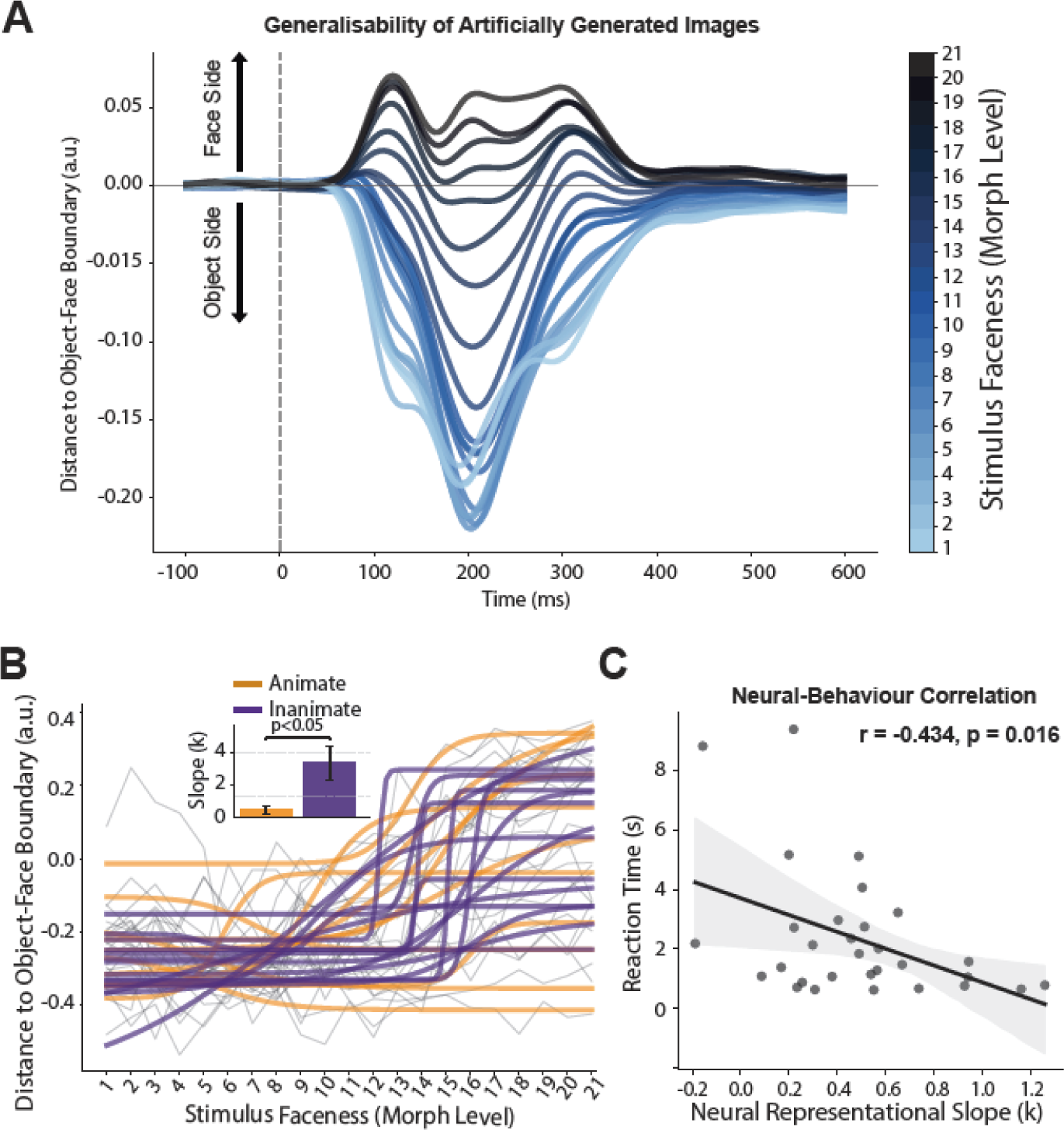
Structure of neural representational space, animacy modulation, and brain–behaviour correlation. **(A) Projection of AI-generated morphs onto the naturalistic object–face decision boundary.** Distance-to-boundary trajectories across time (-100 ms to 600 ms) for all 21 morph levels, derived from multivariate pattern classifiers trained on independent EEG responses to naturalistic object and face stimuli. Positive distances indicate neural representation on the face side of the decision boundary, whereas negative distances reflect projection onto the object side. Maximum representational differentiation between faces and objects occurred at ∼200 ms post-stimulus. **(B) Categorical sharpness across animate vs. inanimate stimulus continua.** Sigmoid functions fitted to decision-boundary distances (within a 50 ms window centered at 200 ms) across morph levels for individual stimulus continua, split by animacy (Animate: orange; Inanimate: purple). Inset bar plot depicts mean (±SE) fitted slope values (*k*), showing significantly steeper slopes for inanimate relative to animate morph continua. **(C) Brain–behaviour correlation between neural category sharpness and task performance.** Scatter plot illustrating the relationship between participants’ fitted neural Sigmoid slopes (*k*) and their median reaction times (s) on forced-choice gender probe trials.

Object-like stimuli were correctly projected onto the object side of the decision boundary, whereas Face-like stimuli mapped onto the face side, with Ambiguous Pareidolic stimuli occupying intermediate positions near the boundary. Face-like and Object-like stimuli also exhibited distinct temporal dynamics. Specifically, face-like images showed an early peak at around 120 ms followed by a sustained representation until approximately 350 ms, whereas Object-like images displayed a single dominant peak at around 200 ms. The maximum representational distance between these extremes occurred at approximately 200 ms, which served as the temporal reference for subsequent analyses. Together, these results demonstrate that our generative AI morph continuum elicited neural representations that are highly consistent with naturalistic object and face stimuli reported in prior literature.

Because the objects depicted in the morphed stimuli varied in semantic categorical content (animate versus inanimate), we hypothesised that there would be systematic variation in categorical thresholds across the stimulus set (i.e., the specific morph level at which an object–face morph begins to be perceived as a face). To test this, we evaluated the slope of the Sigmoid functions fitted to each object–face morph continuum individually within a 50 ms window centered at 200 ms (Figure 6B). Because the base objects that were used to morph into faces belonged to distinct semantic categories, we grouped the 26 morph continua into the evolutionarily fundamental and well-studied inanimate versus animate categories (Mur et al., 2013; Thorat et al., 2019). This partition assigned the top and bottom rows of stimuli in Figure 1B to the inanimate category and the middle rows to the animate category. We then fitted Sigmoid functions to their decision-boundary distances and measured the resulting slopes (Figure 6B).

The slopes for the inanimate category were significantly steeper than those for the animate category (p < 0.05; two-sample t-test), indicating a sharper categorical boundary between inanimate objects and faces than between animate objects and faces. This sharper transition likely reflects the fact that animate objects share animacy features with faces, rendering them closer in neural representational space (and thus less discriminable) relative to inanimate objects and faces, which share no animacy features. These results—demonstrating that animacy organisation structures neural representations in a manner consistent with a broad body of research (Mur et al., 2013; Thorat et al., 2019)—further validate the generalisability of our AI-generated stimulus set and empirical findings.

In line with previous studies (Zhu and Meng, 2019), the behavioural data (Figure 2) exhibited large inter-individual variability in face perception sensitivity, with the greatest variability occurring at the morph level at which individuals first reported perceiving a face within the morphed stimuli. Corresponding variability was present in the Sigmoid functions fitted to their boundary-distance neural data (slope = 0.619 ± 0.212, Figure S3). To establish a direct link between these participant-specific neural sensitivities and behaviour (Rekow et al., 2022), we correlated each participant’s median reaction time (RT) on forced-choice probe trials against their individual neural Sigmoid slope values (Figure 6C).

Participants performed with high accuracy when identifying face gender on probe images/trials with known genders (Accuracy = 97.5 ± 1.54%; Reaction time = 2.34 ± 0.405 s; Mean ± SE). Neural–behavioural correlation analysis revealed a significant negative relationship (*r* = −0.434, *p* = 0.016; Pearson correlation). There was evidence that participants who exhibited sharper, steeper Sigmoidal transition curves in their neural representations resolved the behavioural face identity task significantly faster. This establishes a direct connection between an individual’s neural category boundary sensitivity and their behavioural performance. Furthermore, because the probe trials measured an orthogonal, task-irrelevant feature (face gender) relative to the passive EEG task, this finding suggests that individual categorical perception sensitivity (threshold) is not restricted to object-face and pareidolia perception, but may represent a generalisable feature of visual categorisation.

## Discussion

The goal of this study was to characterise the neural mechanisms underlying face pareidolia by tracking representational dynamics of human brain activity across a continuous, AI-generated 21-level object-to-face morph spectrum. The results reveal a clear spatiotemporal dissociation: early posterior visual processing maintains a linear representation of low-level structural changes in the object-face continuum, whereas delayed frontal areas collapse this physical continuum into a nonlinear, sigmoidal profile that reflects nonlinear categorical representation. Critically, by evaluating directed representational flows, we found that this transformation relies on an initial feed-forward sweep followed by delayed feedback flows. This top-down feedback is preferentially recruited to resolve perceptual ambiguity when sensory input is highly ambiguous near the category boundary, which is occupied by pareidolic stimuli. These findings demonstrate that face pareidolia relies on recurrent processing, reconciling prior conflicting models that emphasised either strictly bottom-up, feed-forward mechanisms or predominantly top-down modulation.

### Dynamic Recurrence Unifies Feedforward and Top-Down Frameworks

The spatiotemporal cascade observed in our dataset strongly aligns with and extends recent findings in face pareidolia (Wardle et al., 2020; Robinson et al., 2025). Using EEG and functional magnetic resonance imaging (fMRI), that work established that stimuli evoking face pareidolia engage the same ventrotemporal neural circuitry and trigger early face-selective electrophysiological responses (such as the N170) as typical human faces supporting similar findings (Hadjikhani et al., 2009; Rekow et al., 2022; Cerrahoğlu et al., 2025). Crucially, Wardle et al. (2020) and Robinson et al. (2025) identified a key temporal transition: despite the rapid, spontaneous emergence of face-like representations (90–130 ms), the underlying neural profile shifts toward an object-like representation at longer latencies (150–210 ms), suggesting that the brain generates multiple distinct representations for pareidolic inputs. This dual-stage phenomenon aligns with other neurophysiological evidence showing that early ventral visual areas automatically extract structural facial features before late-stage processing stabilises object identity (Keysers et al., 2001; Moeller et al., 2017). Our results replicate this fundamental temporal evolution while grounding it within a formal mathematical framework. By modelling the full similarity geometry, we have shown that early face-like signals in posterior channels reflect a rapid, continuous tracking of low-level visual features (the Linear model). The critical transition noted by previous studies corresponds directly to our frontal sigmoidal resolution window (180–350 ms). Rather than a simple overwriting of sensory representations or a complete re-tuning of early visual cortex, our data reveal a spatial and temporal architectural split. Specifically, we find that, after 200ms, posterior channels continue to track structural feature changes while anterior networks simultaneously extract categorical identity. This provides quantitative support for the parallel-representation hypothesis (Contini et al., 2017; Kietzmann et al., 2019), demonstrating how the human visual system preserves an object’s physical structure in sensory cortex while concurrently resolving its illusory facial identity in higher-order networks.

While previous literature on face pareidolia has largely pitted feed-forward accounts of automatic sensory extraction against top-down models of expectation-driven perceptual construction, our empirical findings demonstrate that these perspectives capture complementary axes of a unified computational loop. Crucially, these two paradigms have historically evaluated pareidolia at distinct representational levels: feed-forward models assess continuous visual feature extraction, whereas top-down frameworks evaluate discrete categorical decisions. Studies supporting feed-forward accounts suggest that pareidolic stimuli share early image-level and structural representational profiles with natural faces, distinguishing them from standard non-face objects in posterior visual regions (Churches et al., 2009; Palmer & Clifford, 2020; Rekow et al., 2022; Cerrahoğlu et al., 2025). Conversely, studies supporting top-down accounts focus primarily on categorical decision-making under high perceptual uncertainty, demonstrating that higher-order expectations and frontoparietal feedback force ambiguous sensory inputs into discrete, binary identity categories (Bar et al., 2006; Liu et al., 2014; Lu et al., 2017; Powers et al., 2016). By mapping directed representational flows across a continuous morph spectrum, our results unify these findings within a single, multi-stage architecture. Early posterior processing operates via a rapid feed-forward sweep that continuously tracks structural feature gradients without committing to a categorical decision (Felleman & Van Essen, 1991; Riesenhuber & Poggio, 1999; Kravitz et al., 2013). This feed-forward input provides the raw visual evidence that is subsequently transformed into a discrete, sigmoidal categorical boundary within prefrontal networks (Freedman et al., 2001; Miller & Cohen, 2001; Goldstone & Hendrickson, 2010). When sensory ambiguity peaks near this boundary—the precise ‘zone of uncertainty’ occupied by pareidolic stimuli—these anterior representations initiate sustained top-down feedback loops to early visual regions to resolve perceptual uncertainty (Hochstein & Ahissar, 2002; Lamme & Roelfsema, 2000). Rather than treating pareidolia as a low-level sensory phenomenon (Churches et al., 2009, 2014) or an isolated top-down hallucination (Powers et al., 2016; Partos et al., 2016), our findings establish it as the emergent product of a feedforward-initiated, top-down-resolved recurrent loop.

### Theoretical Implications for Predictive Coding Frameworks

The selective recruitment of top-down feedback for highly ambiguous or pareidolic stimuli provides strong empirical support for predictive coding frameworks of visual processing (Friston, 2005; Summerfield & de Lange, 2014). When a sensory signal contains conflicting features—straddling the boundary between an everyday object and a human face—the bottom-up feedforward sweep generates a substantial prediction error in frontoparietal networks. To resolve this uncertainty, anterior “initial guesses” are projected back to early sensory areas via feedback loops to dynamically optimise local representations (Bar et al., 2006; Mechelli et al., 2004; Lu et al., 2017). This mechanistic architecture provides a plausible neural basis for recent behavioural evidence showing that pareidolic faces are not only detected rapidly, but also engage the same downstream, high-level social processing mechanisms as real faces (such as sensory adaptation and gaze-direction coding; Palmer & Clifford, 2020). Top-down feedback allows the brain to rapidly commit to a face interpretation, enabling the immediate extraction of social information from inanimate objects.

### Divergence Between Biological Recurrence and Artificial Feedforward Architectures

Recent computational work has shown that feedforward deep convolutional neural networks (CNNs) optimised for dual tasks, such as face identification and object categorisation, can spontaneously represent illusory faces (Gupta & Dobs, 2025; Hamilton et al., 2024). However, this feedforward emergence reveals key limitations. While CNNs detect coarse face-like configurations, they fail to capture the flexible, context-dependent ambiguity resolution seen in human behavior (Hamilton et al., 2024). Furthermore, representational similarities in AI do not mean artificial models utilise the same underlying biological mechanisms (Linsley et al., 2025). While feedforward artificial networks can extract structural face-like primitives from inanimate objects, our findings demonstrate a key divergence in human visual architecture. The human brain does not rely solely on feedforward feature extraction to dictate perception, but instead deploys an anterior-to-posterior recurrent loop to resolve categorical uncertainty (Karimi-Rouzbahani et al., 2021; Abouhadi & Karimi-Rouzbahani, 2026). Purely feedforward sweeps are insufficient to capture temporal representational dynamics, which require recurrent dynamics—both local lateral and top-down feedback loops—to refine category boundaries over time (Kietzmann et al., 2019). Incorporating dynamic, top-down feedback mechanisms into artificial vision architectures may therefore be essential for capturing the flexible, uncertainty-resolving dynamics that characterise biological perception.

### Parametric Ambiguity Resolution Across Generative Continua

A key advantage of the present paradigm lies in its ability to map the mathematical profile of the perceptual transition zone across a continuous, parametric continuum. Prior face pareidolia paradigms have yielded fundamental insights by contrasting distinct categories (such as human faces, pareidolic objects, and semantically matched control objects e.g., Wardle et al., 2020; Rekow et al., 2022) or by embedding faces within varying levels of visual noise (e.g., Liu et al., 2014; Ales et al., 2012; Karimi-Rouzbahani et al., 2021). Real-world pareidolic experiences naturally span a continuous structural spectrum, however, ranging from subtle, vague facial hints to striking, unambiguous face-like configurations. By leveraging an AI text-to-image diffusion pipeline to generate a continuous 21-level object-to-face spectrum, our methodology extends beyond binary classification models and noise-degradation frameworks to capture how the brain resolves perceptual ambiguity along a smooth visual continuum. Our approach has three major advantages. First, the continuous steps allowed us to adjudicate between competing psychometric and neural hypotheses (i.e., Linear, Sigmoid and Step), characterising the spatiotemporal representation of each prediction in the brain. Second, we extracted individual and stimulus-specific transition parameters (slopes), revealing that animacy priors structurally affect the category boundary, and that individual neural sigmoidal steepness directly predicts behavioural response speed on probe trials. Finally, we investigated and compared the highly conflicted, ambiguous pareidolic transition zone (Levels 8–14 in our morphing contiuum) against the unambiguous ends of the spectrum. This division confirmed that while unambiguous objects and faces are resolved efficiently via feedforward sweeps, ambiguous (pareidolic) images selectively recruit a prolonged top-down prefrontal feedback loop (Kar & DiCarlo, 2021; Kreiman & Serre, 2020; Karimi-Rouzbahani, 2026; Tang et al., 2018).

### Methodological Advances in Transformed Representational Flow and High-Temporal Constraints

Methodologically, uncovering these spatiotemporal dynamics required moving beyond traditional functional connectivity methods that rely on univariate correlated activities and are blind to the content of transferred representations (Anzellotti & Coutanche, 2018; Karimi-Rouzbahani et al., 2022). While next-generation connectivity analyses focus on information transfer (Karimi-Rouzbahani et al., 2022; Lorenz et al., 2025), these approaches still miss critical signals if information undergoes computational transformation during transfer. The transformation-based RCA framework developed here yields two key insights that would have been missed using even the latest untransformed RCA methods (Abouhadi & Karimi-Rouzbahani, 2026). First, it captured transformed representational flows from linear profiles in posterior areas to sigmoidal representations in anterior areas, which we showed would have been missed if using untransformed RCA (Figure S1). Second, by systematically stripping away historical representations in the destination region, our approach ensured that measured sigmoidal representations in anterior areas reflected true transformed representational flow from posterior areas rather than parallel, uncoupled responses to visual onset.

Our paradigm also suggests that the brain extracts low-level structural features and initiates top-down recurrent loops under ultra-rapid, 50-ms visual presentation constraints within an RSVP stream. While many of previous electrophysiological, fMRI, and behavioural investigations of face pareidolia have relied on longer or extended visual exposures (e.g., 200–500 ms) (Churches et al., 2009, 2014; Akdeniz, 2020; Palmer & Clifford, 2020), there is now evidence that face pareidolia can be studied with shorter stimulation durations (Rekow et al., 2022). Demonstrating that this feedforward-to-feedback resolution operates so robustly under such stringent temporal limits underscores the remarkable efficiency and automaticity of the anterior-posterior interactions in processing illusory social primitives (Philiastides & Sajda, 2006; DiCarlo et al., 2012). Future research utilising high-density intracranial recordings (iEEG) or combined MEG-fMRI could further refine this spatiotemporal map, tracking how recurrent loops operate across localised cortical layers to adjudicate between competing perceptual interpretations.

## Acknowledgements

HKR was supported by an Australian Research Council DECRA Fellowship DE230100608. JBM was supported by a National Health and Medical Research Council Investigator Grant (GNT2010141) and the Canadian Institute for Advanced Research (CIFAR). We would like to thank Mr Mohammad Hossein Niki-Maleki for generating the imageset and collecting the behavioural dataset and Mr Douglas Cribb for collecting the EEG dataset. We also thank Drs Jessica Taubert and Amanda K. Robinson for providing insightful feedback on the project.

## Code and Data Availability

The analysis code is available at: https://github.com/HamidKarimi-Rouzbahani/Face_pareidolia_morphed_stims_EEG.git. The EEG and behavioural data is available at: https://osf.io/r9sqe/overview?view_only=8a8bf87f4ef847bc95521511dd01cd46

## Authors Contributions

**Conceptualization**: HKR; **Methodology**: HKR & JBM; **Software**: HKR; **Validation**: HKR; **Formal Analysis**: HKR; **Investigation**: HKR; **Resources**: HKR & JBM; **Data Curation**: HKR; **Writing – Original Draft**: HKR; **Writing – Review & Editing**: HKR & JBM; **Visualization**: HKR; **Supervision**: JBM; **Project Administration**: HKR; **Funding Acquisition**: HKR & JBM.

## Supplementary materials

**Text S1** The steps taken in implementing RCA are explained in detail below:

### 1. Instantaneous Representation Amplitude (Partial Correlation Scalar)

To isolate unique model-specific representations in a given brain region at time *t*, the target model RDM vector **m**_target_ and neural RDM vector **r**_neural_(*t*) are residualised against nuisance model RDMs **M**_nuisance_:

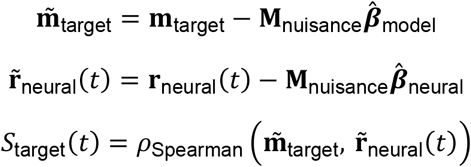

where:

- **m**_target_ is the upper-triangular vector of the target model RDM (e.g., Linear, Sigmoid, or Step).
- **r**_neural_(*t*) is the upper-triangular vector of the empirical neural RDM calculated across conditions at time *t*.
- **M**_nuisance_ is a matrix containing the upper-triangular vectors of the remaining, non-target hypothesis model RDMs (e.g., when target is Linear, nuisance models are Sigmoid and Step).
- 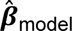 is the vector of Ordinary Least Squares (OLS) regression coefficients fitting **M**_nuisance_ to **m**_target_ (i.e., 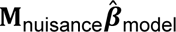 represents the nuisance variance predicted within the target model).
- 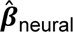 is the vector of OLS regression coefficients fitting **M**_nuisance_ to **r**_neural_(*t*) (i.e., 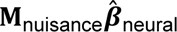 represents the nuisance model variance predicted within the neural RDM).
- 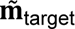 and 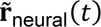 are the history- and nuisance-cleared residual vectors for the target model and neural data, respectively.
- *ρ*_Spearman_ is Spearman’s rank correlation coefficient.
- *S*_target_(*t*) is the resulting partial correlation scalar reflecting unique representational coding for the target model at time *t*.

### 2. Destination History-Stripping (Residualisation)

To isolate *novel* updates transferred to the destination region at time *t*, the destination region’s past neural state **r**_dest_(*t* − Δ*t*) is regressed out of both the current destination neural RDM **r**_dest_(*t*) and the current target model RDM **m**_target_:

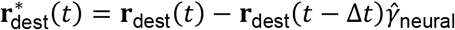

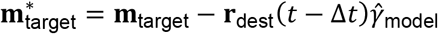

where:

- Δ*t* is the inter-regional transmission temporal delay (evaluated from 5 ms to 200 ms).
- **r**_dest_(*t* − Δ*t*) is the upper-triangular vector of the destination region’s neural RDM at the preceding historical time point *t* − Δ*t*.
- **r**_dest_(*t*) is the upper-triangular vector of the destination region’s neural RDM at the current time point *t*.
- 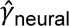 is the OLS regression slope estimate of **r**_dest_(*t* − Δ*t*) predicting **r**_dest_(*t*).
- 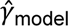 is the OLS regression slope estimate of **r**_dest_(*t* − Δ*t*) predicting **m**_target_.
- 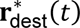 and 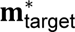 are the history-stripped residuals representing novel variance in the current neural activity and model template, respectively.
- 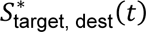 is the history-stripped scalar update computed via partial correlation using these residual vectors 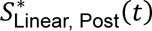 and 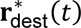 while controlling for nuisance models.

### 3. Representational Directed Flow (Scalar Product)

The directional informational flow across time step *t* and inter-regional delay Δ*t* is defined as the scalar product of the historical source representation amplitude and the history-stripped destination residual update:

**Transformed Directed Flow (Linear** ↔ **Sigmoid)**

- Feedforward Transformed Flow **(FF_trans_):**

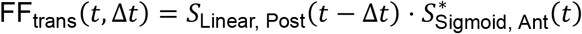

- **Feedback Transformed Flow** (FB_trans_):

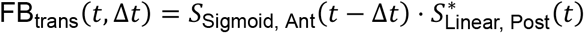

**Non-Transformed Flow Controls (Identical Mapping *M*** → ***M*)**

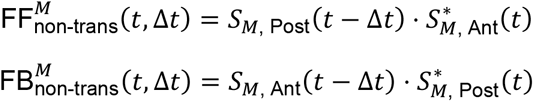

where:

- *S*_Linear, Post_(*t* − Δ*t*) is the instantaneous Linear partial correlation scalar in Posterior channels at historical time *t* − Δ*t*.
- 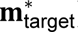 is the history-stripped Sigmoidal partial correlation scalar update in Anterior channels at current time *t*.
- *S*_Sigmoid, Ant_(*t* − Δ*t*) is the instantaneous Sigmoidal partial correlation scalar in Anterior channels at historical time *t* − Δ*t*.
- 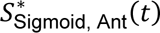 is the history-stripped Linear partial correlation scalar update in Posterior channels at current time *t*.
- *M* represents the non-transformed structural pathway model, where *M* ∈ {Linear,Sigmoid,Step}.

---

**Figure S1.**
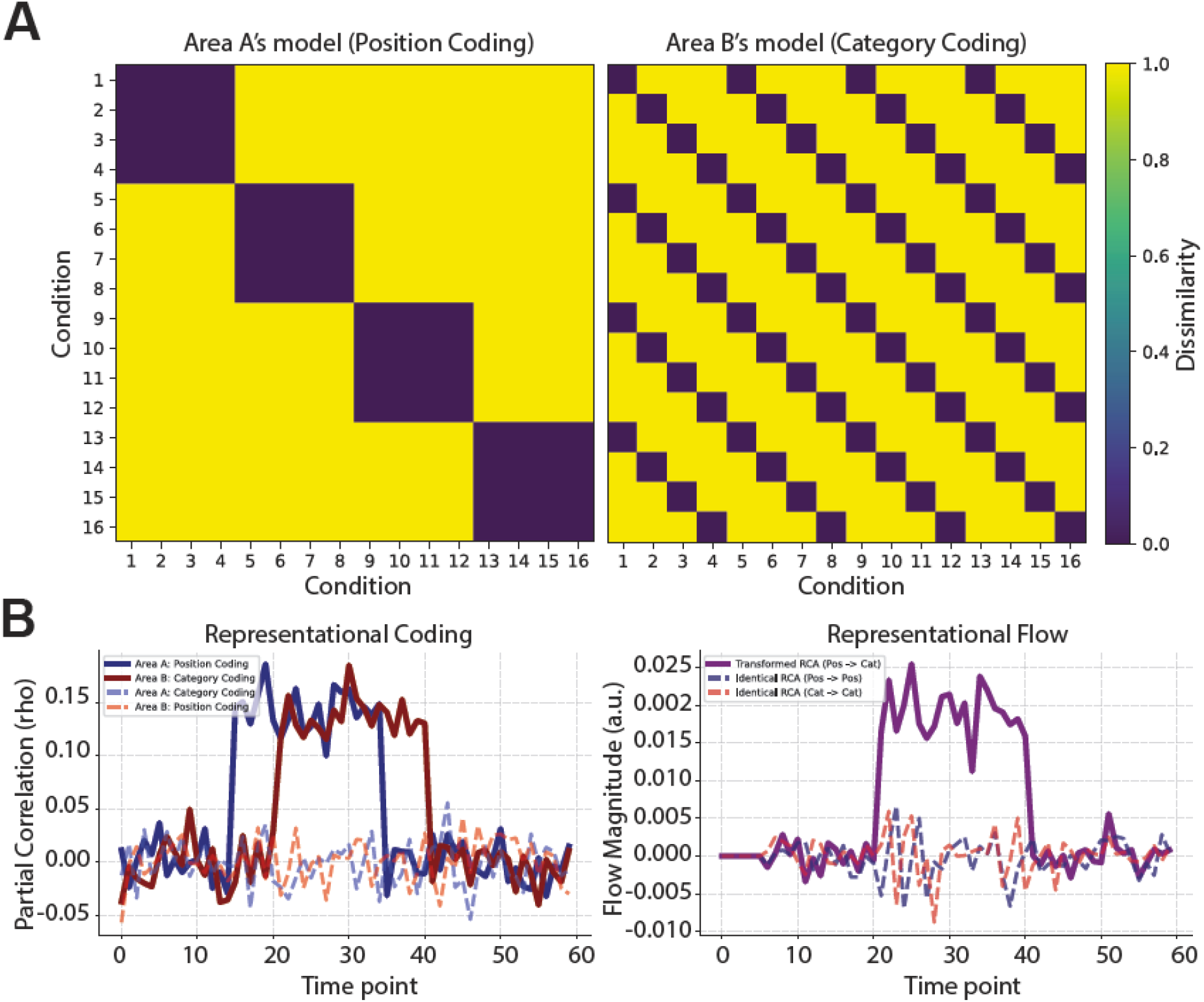
Simulation of cross-regional representational transformation and its selective detection via transformation RCA. (A) Ground-truth RDM model templates. RDM model templates across a 16-condition factorial design. *Left:* Area A’s model RDM captures *spatial position* coding, structured as four 4 × 4 block clusters along the diagonal. *Right:* Area B’s model RDM captures orthogonal *category* coding, structured as a periodic tiled pattern across the 16 conditions. Colour bar indicates normalised Hamming dissimilarity (0.0 to 1.0). **(B) Time-resolved information coding and represenational flow.** Time-resolved metrics evaluated across 60 simulated time points (*N* = 20 subjects; activation window = 20 time points; true inter-regional transmission delay *τ* = 5 time points). *Left (Representational Coding):* Time-resolved partial Spearman correlation profiles confirm that Area A selectively encodes *spatial position* during the initial activation window (solid dark blue line; time points 15–35), whereas Area B selectively encodes *category* information following the simulated transmission delay (solid dark red line; time points 20–40). Cross-feature/untransformed coding traces in both regions remain at noise baseline (dashed lines). *Right (Reprsentational Flow):* Time-resolved RCA flow evaluated at the simulated delay (*τ* = 5). Transformed RCA (Pos → Cat; solid purple line)—computed by evaluating the cross-regional mapping between Area A’s past position-coding scalar at *t* − *τ* and Area B’s current category-coding scalar at *t*—successfully isolates a robust peak in directed information transfer corresponding to the activation window. In contrast, identical/untransformed RCA control pathways (Pos → Pos, dashed navy line; Cat → Cat, dashed red line) remain around baseline throughout the epoch, demonstrating that conventional model-based RCA fails to detect inter-regional flow when representational content undergoes transformation across areas.

**Figure S2.**
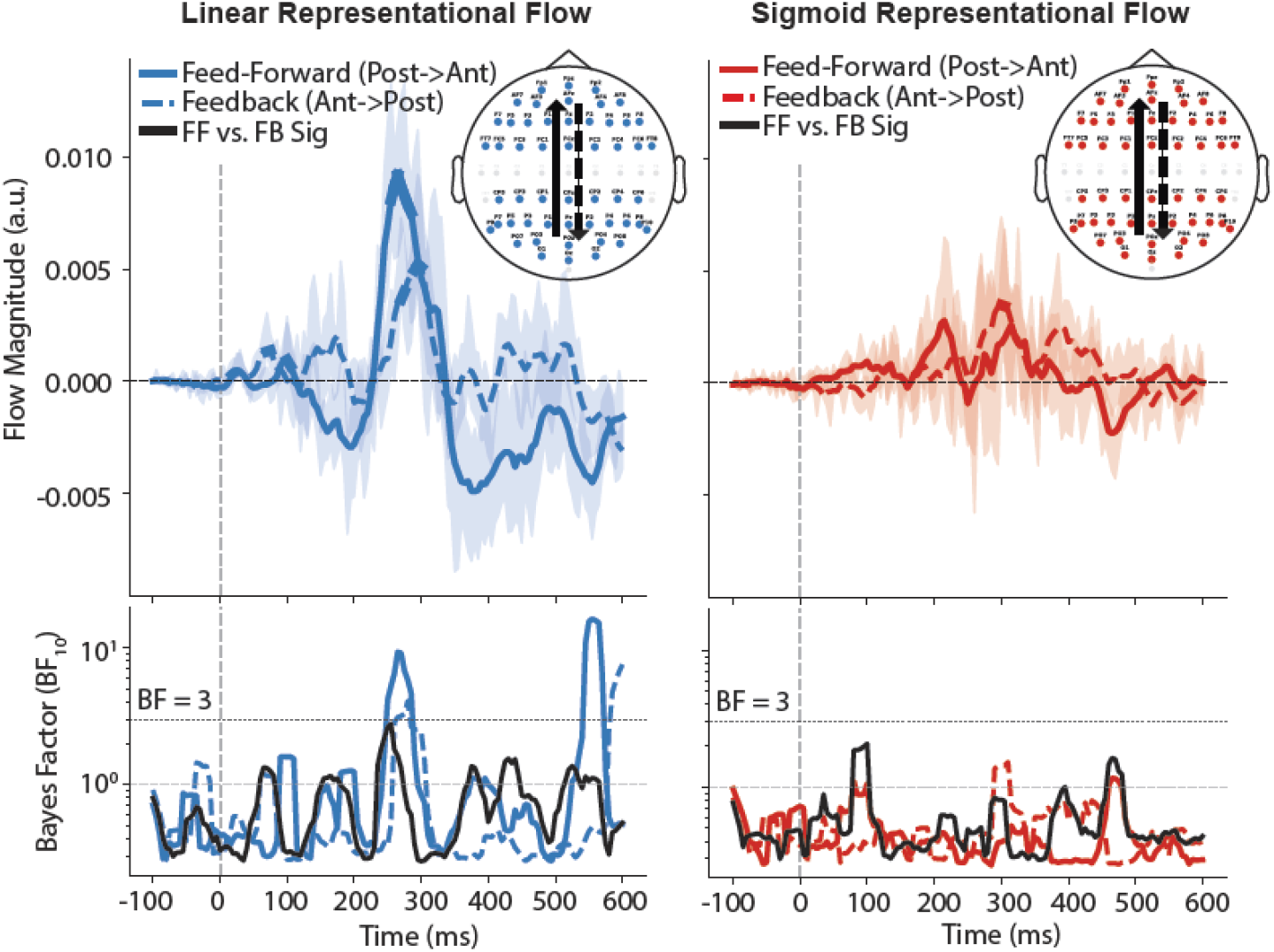
Non-transformed representational connectivity. Inter-areal representational connectivity tracking communication purely within un-transformed models across source and destination areas for Linear (left panels) and Sigmoid (right panels) profiles. Feed-forward flow (posterior-to-anterior; solid lines) and feedback flow (anterior-to-posterior; dashed lines) magnitudes are plotted in the upper panels alongside difference trajectories (black lines), with corresponding Bayes Factors (BF_10_) depicted in the lower panels. Horizontal black bars indicate time points when there was evidence (BF > 3) for higher feed-forward than feedback flows.

**Figure S3.**
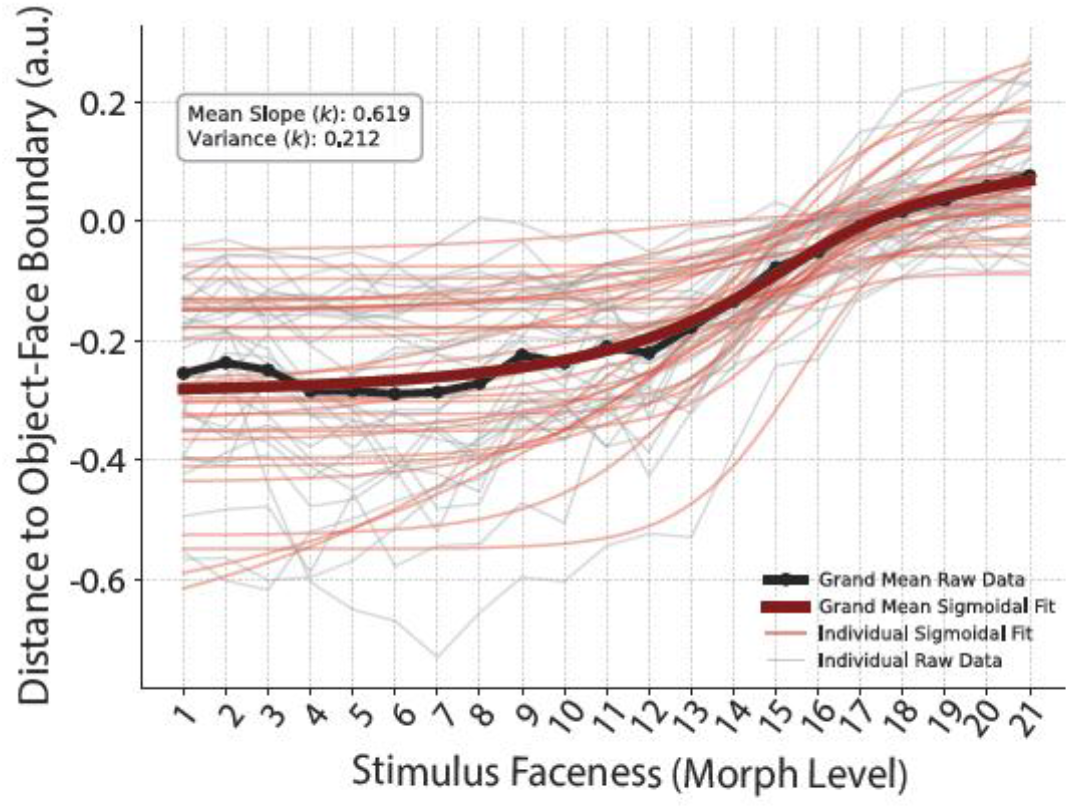
Quantification of perceptual categorisation dynamics across stimulus morph levels via sigmoidal modelling. Individual participant and grand-mean neural representational distances plotted relative to the object–face decision boundary at 200ms across 21 morph levels. Individual participant raw distance metrics (thin grey lines) were fitted with logistic Sigmoidal functions (thin light-red lines) to quantify individual nonlinear transition profiles across the continuum. The black line with filled circles denotes the grand-mean empirical raw data across subjects, whereas the thick dark-red curve highlights the fitted grand-mean Sigmoidal response profile. The inset box reports group-level Sigmoidal slope statistics—reflecting individual variability in categorical boundary steepness—with a mean slope coefficient (*k* = 0.619) and cross-subject slope variance (*k* = 0.212).

## Notes

### Competing Interest Statement

The authors have declared no competing interest.

